# An interaction between β’-COP and the ArfGAP, Glo3, maintains post-Golgi cargo recycling

**DOI:** 10.1101/2022.05.25.493481

**Authors:** Boyang Xie, Clara Guillem, Christian Jung, Amy K. Kendall, Swapneeta Date, Jordan T. Best, Todd R. Graham, Lauren P. Jackson

## Abstract

The essential COPI vesicular coat mediates retrieval of key transmembrane proteins at the Golgi and endosomes following recruitment by the small GTPase, Arf1. ArfGAP proteins regulate COPI coats, but molecular details for COPI recognition by ArfGAP proteins remain elusive. Biochemical and biophysical data reveal how β’-COP propeller domains directly engage the yeast ArfGAP, Glo3, with a low micromolar binding affinity (K_D_ ~1 µM). Calorimetry data demonstrate both β’-COP propeller domains are required to bind Glo3 using electrostatic interactions. An acidic patch on β’-COP (D437/D450) interacts with Glo3 lysine residues located within the BoCCS (Binding of Coatomer, Cargo, and SNAREs) region. Targeted point mutations in either Glo3 BoCCS or β’-COP abrogate the interaction *in vitro*, and loss of the β’-COP/Glo3 interaction drives Ste2 mis-sorting to the vacuole and aberrant Golgi morphology in budding yeast. Together, these data suggest cells require the β’-COP/Glo3 interaction for cargo recycling via endosomes and the TGN, where β’-COP may serve as a molecular platform to coordinate binding to multiple protein partners, including Glo3, Arf1, and the COPI F-subcomplex.

## Introduction

The highly conserved COPI coat^1–3^ is essential for vesicular membrane trafficking in eukaryotes. COPI has many established roles, including retrieval of endoplasmic reticulum (ER) proteins from the Golgi back to the ER; cycling of proteins between the ER and Golgi; retrograde^4^ and anterograde^5^ trafficking within the Golgi stack; and from endosomes to the TGN^6^. COPI has been implicated in lipid homeostasis^7,8^, viral replication^9^, and pathogen entry^10–12^. Mutations or upregulation of COPI subunits have been have specifically linked to both inherited and acquired diseases, including microcephaly^13^ and cancer^14^. Viral glycoproteins use established COPI motifs to circumvent host immunity.^15,16^

The COPI heptamer (α−/β−/β’-/γ−/δ−/ε−/ζ−COP subunits) is recruited *en bloc* onto Golgi membranes^17^ following recruitment by the small GTPase, Arf1, and cargo. COPI historically has been conceptually divided into the B-subcomplex (α/β’/ε) and F-subcomplex (β/δ/γ/ζ). The F-subcomplex is both structurally and functionally related to the AP clathrin adaptor complexes^18,19^. The B-subcomplex is often compared to clathrin, but it shares no evolutionary history^18^. Structural data demonstrate COPI is an interwoven coat with multiple contacts between subunits^20–22^, in contrast to the layered structure adopted by clathrin^23^ and COPII^24,25^ coats. The COPI B-subcomplex contains four WD-repeat (also known as β-propeller) domains, which are interaction platforms that can accommodate multiple protein binding partners. The α− and β’-COP N-terminal WD-repeat domains bind dilysine motifs in transmembrane cargoes^26,27^ and K63-linked ubiquitin chains^6^. The COPI WD-repeat domains found in α− and β’-COP are strong candidates for recognizing other proteins, including regulators.

ArfGAP proteins are critical regulators of COPI function, but their precise molecular roles remain poorly understood. ArfGAPs have been implicated in coat assembly^28,29^; cargo/SNARE sorting^30–32^; and coat disassembly^33^. Yeast contain two essential ArfGAP proteins, Glo3 (Figure 1A) and Gcs1, which have overlapping functions^34^. Both ArfGAPs have homologs/orthologs in mammalians cells: Gcs1 corresponds to ArfGAP1, while Glo3 corresponds to ArfGAP2/3. Budding yeast tolerate deletion of either *GLO3* or *GCS1* individually but deletion of both genes is lethal^35^. Glo3 GAP activity is an essential function^29,36^. Strains harboring the GAP-dead version of Glo3 (R59K) are not viable, even in the presence of Gcs1, which implies a dominant role for Glo3. In mammalian cultured cell lines, mutations in key basic residues of ArfGAP2/3 have been shown to affect its localization and function.^37^

**Figure 1.**
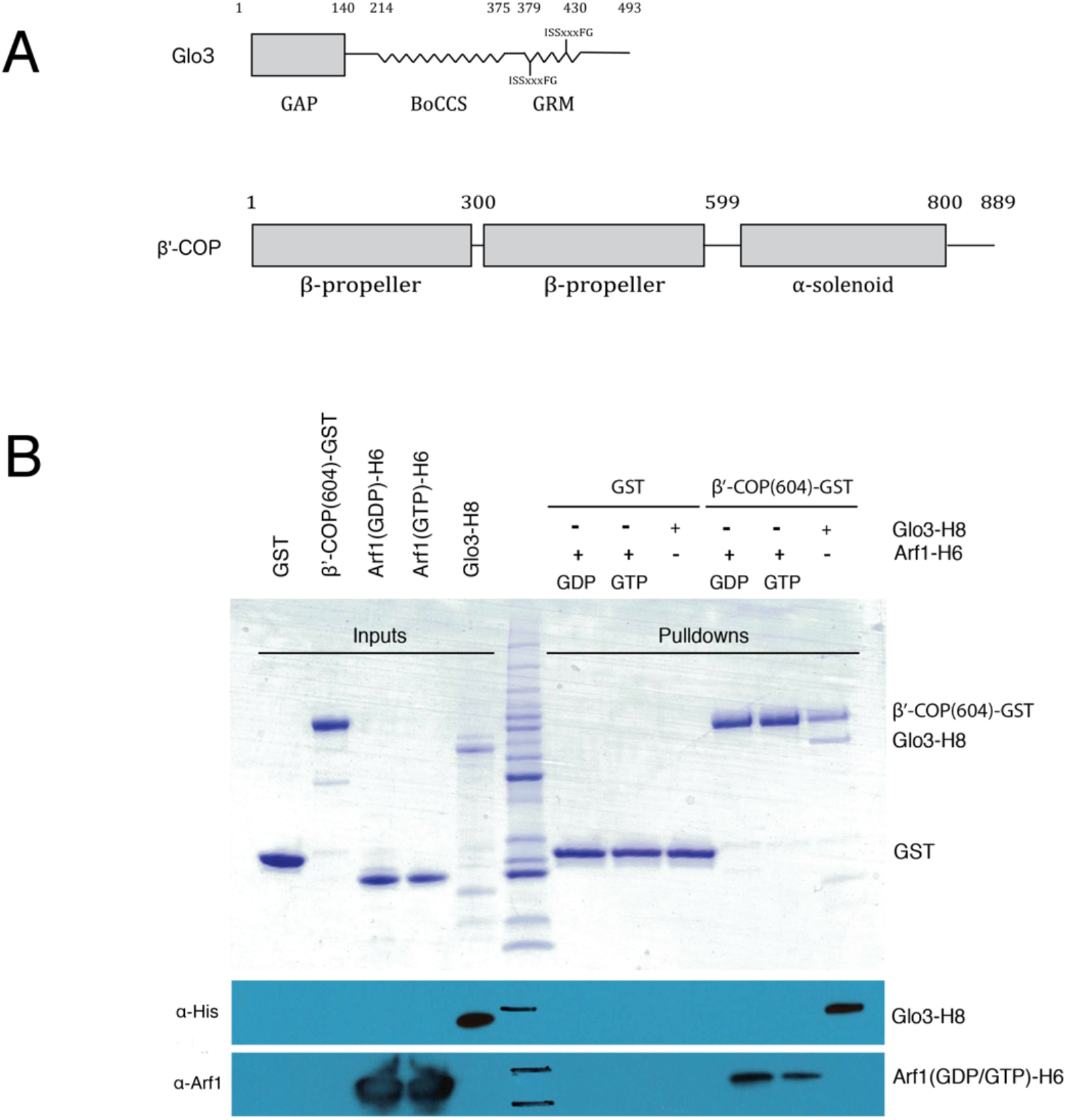
β’-COP propeller (WD-repeat) domains directly bind Glo3 and Arf1 *in vitro*. (A) Schematics of yeast Glo3 and β’-COP proteins. Glo3 contains a GAP domain (residues 1-140), BoCCS region (defined as residues 214-375), and GRM region (residues 375-493). β’-COP contains two WD-repeat (also known as β-propeller) domains followed by an α-solenoid. The N-terminal propeller domain binds dilysine motifs in transmembrane cargo, while the α-solenoid interacts with α-COP. (B) Pulldown experiments using GST-tagged β’-COP propeller domains (residues 1-604 with C-terminal GST tag) as bait and either full-length Arf1-H6 or full-length Glo3-H8 as prey. We tested binding to both GDP-locked (T31N) Arf1 and GTP-locked (Q71L) Arf1. β’-COP interacts directly with both Arf1 constructs and with Glo3 *in vitro*. The top panel shows a Coomassie-stained SDS-PAGE gel, while the bottom two panels show Western blots against the Glo3 His8 tag (α-His; Abcam NB100-63173) or against yeast Arf1 (α-Arf1, Todd Graham lab, Vanderbilt University).

There are very limited molecular data indicating how ArfGAPs interact with the COPI coat and the precise roles they play in trafficking. Classically, ArfGAPs were thought to drive vesicle uncoating by hydrolyzing GTP and releasing Arf1(GTP) from membranes; in this model, the ArfGAP serves to recycle COPI. However, Glo3 has been shown to play important roles in coat assembly, including cargo selection and SNARE binding via the BoCCS region in Glo3^32^. Biochemical data from cell lysates^36^ suggest Glo3, but not Gcs1, stably associates with COPI. Both genetic^38,39^ and affinity-capture data^40^ in budding yeast suggest an interaction between COPI and Glo3, while the ArfGAP2/3 GAP domain has been visualized in a cryo-electron tomography (cryo-ET) reconstructions^21^ near the B-subcomplex. In yeast cell lysates, Glo3 interacts with γ-COP appendage domain^41^ and associates stably with COPI^36^. Despite multiple lines of evidence, the biochemical basis for a direct interaction between COPI subunits and Glo3 has not been established.

Previous work^26^ and data on other coat proteins^42–44^ led to the hypothesis that β’-COP functions as a molecular platform to engage multiple protein partners within the COPI coat. In this work, we set out to identify additional binding partners for the β’-COP WD-repeat (also known as propeller) domains. We identified the ArfGAP, Glo3, and Arf1 as direct β’-COP binding partners *in vitro*. We characterized the β’-COP/Glo3 interaction by mapping specific residues on both proteins; quantifying binding affinities; and testing structure-based mutants *in vitro*. We tested whether disrupting the β’-COP/Glo3 interaction affected dilysine cargo binding *in vitro*, as well as whether loss of the interaction affected cargo sorting and Golgi morphology in budding yeast. Together, these new data identify an expanded role for the β’-COP subunit and suggest a compelling model for the β’-COP/Glo3/Arf1 complex on Golgi membranes.

## Results

### β’-COP directly binds Glo3 and Arf1 *in vitro*

WD-repeat (also known as β-propeller) domains are known protein binding platforms^45^. In preliminary work, we sought to identify other direct binding partners for the β’-COP propeller domains, beyond the well-established dilysine motif cargo^26,27^. We used recombinant purified β’-COP protein (residues 1-604 with a C-terminal GST tag or β’604-GST; Figure 1A) as bait in pulldown experiments from budding yeast cell lysates and identified both Glo3 and Arf1 as potential binding partners (Figure S1A). We next undertook biochemical pulldown experiments to ascertain whether purified recombinant proteins directly bind each other (Figure 1B). The data indicate β’604-GST binds directly to both Glo3 and Arf1 in either nucleotide-bound state (Figure 1B; Figure S1B). The β’-COP/Glo3 interaction can be visualized with Coomassie staining, which suggested the binding affinity (K_D_) between β’-COP and Glo3 lies in the low micromolar range. A three-way pulldown indicated all three proteins interact simultaneously with each other (Figure S1B). When five-fold molar excess Arf1 is added, the β’604-GST/Glo3 complex does not appear to favor either nucleotide bound state (GDP-locked T31N or GTP-locked Q71L). These data provide the first biochemical evidence for a direct interaction between β’-COP and Glo3, and between β’-COP and Arf1. These data are further supported by structural data from COPI coats reconstituted *in vitro*, which place β’-COP and Arf1 adjacent to each other in reconstructions^20^.

We next set out to ascertain which part of Glo3 binds β’-COP. Glo3 (Figure 1A) contains a GAP domain; an unstructured middle region called the BoCCS (for <u>B</u>inding of <u>C</u>oatomer, <u>C</u>argo, and <u>S</u>NAREs^32^; and a C-terminal GRM domain (<u>G</u>olgi <u>R</u>egulatory <u>M</u>otif). We purified recombinant GST-fusion proteins encompassing each portion of Glo3 and undertook pulldown experiments (Figure S2). These data indicate only the Glo3 BoCCS region can pull down β’604-H6. The BoCCS region has been previously implicated in binding TAP-purified COPI from yeast cell lysates^36^, but the COPI subunit mediating the interaction was unknown. Our data reveal for the first time how the β’-COP subunit directly binds Glo3 BoCCS.

### Glo3 BoCCS binds both β’-COP propellers with low micromolar affinity

The initial biochemical data indicated the interaction between β’-COP and Glo3 should be strong enough to be quantified using calorimetry. We thus mapped the binding region within Glo3 using isothermal titration calorimetry (ITC) to quantify binding affinities between purified recombinant proteins (Figure 2A; Figure S3; Table S1). All affinity tags were removed from both Glo3 and β’1-604 prior to running calorimetry experiments, and the initial goal was to ascertain the minimal binding fragment for pursuing structural studies. Pulldown experiments suggested a low micromolar interaction between full-length Glo3 and the β’-COP propeller domains (residues 1-604), and only the BoCCS region appeared to interact directly with β’-COP (Figure S2). Initial experiments using the full BoCCS region (residues 208-375) revealed a low micromolar K_D_ (Figure S3A) and 1:1 stoichiometry. We generated a series of Glo3 constructs truncated at either the N-or C-terminal end; we purified each protein and quantified binding to β’604 by ITC (Figure 2A; Figure S3B). Although longer purified Glo3 constructs are unstable over time (Figure S2), shorter Glo3 fragments used in ITC experiments exhibited high purity and stability after purification (representative example SDS-PAGE gel shown in Figure S3C).

**Figure 2.**
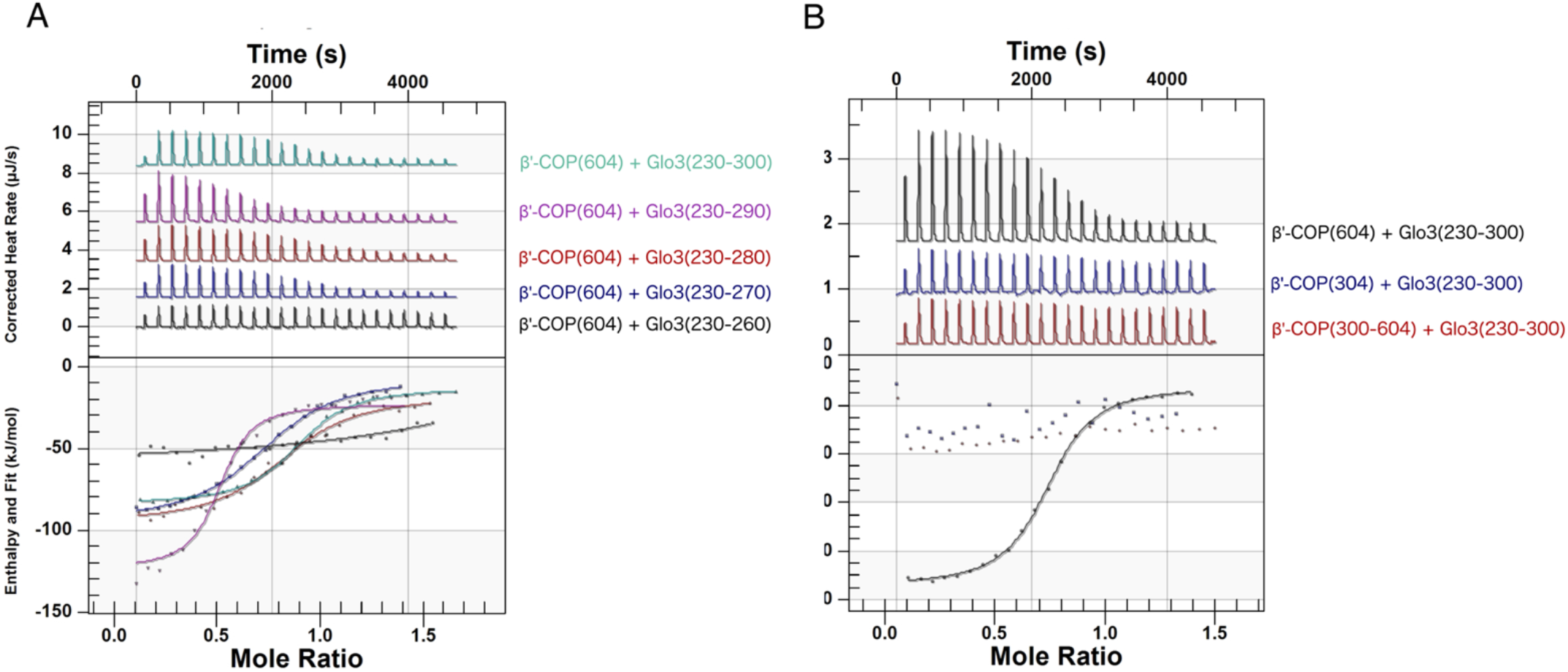
Residues within the Glo3 BoCCS region directly bind both β’-COP propellers with low micromolar affinity. (A) Purified recombinant proteins (untagged β’-COP residues 1-604 and Glo3 fragments as labelled) were used in isothermal titration calorimetry (ITC) experiments to quantify binding affinities; representative traces are shown. β’-COP binds a Glo3 fragment located within the BoCCS region. The highest affinity interaction occurs between β’-COP 1-604 and Glo3 residues 230-290, but all fragments exhibit low micromolar K_D_ values (0.8-6 µM) and 1:1 stoichiometry (Table S1). (B) Representative ITC experiments between untagged Glo3 BoCCS fragment (residues 230-300) and N-terminal β’-COP propeller (residues 1-304), C-terminal β’-COP propeller (residues 300-604), or both β’-COP propellers (residues 1-604). Each propeller domain on its own is insufficient to produce measurable binding by calorimetry, which suggests both propellers are required to bind Glo3.

We observed the strongest interaction between the β’-COP propeller domains and Glo3 residues 230-290 (average K_D_= 0.6±0.2 µM, N=3 runs; representative trace in Figure 2A; Table S1). We saw no difference in binding affinity when comparing β’-COP binding to Glo3 residues 220-290 or Glo3 residues 230-290, but loss of Glo3 residues 230-240 abrogated measurable ITC binding (Figure S3B). Truncations at the C-terminal end resulted in progressively weaker interactions (Figure 2A). These data define Glo3 residues 230-290 as the key region that interacts directly with β’-COP propellers.

We next tested which β’-COP propeller domain is required for the interaction with Glo3 (Figure 2B). Specifically, we tested Glo3 residues 230-300 with each propeller domain on its own (either β’-COP residues 1-304 or residues 305-604) and with both propeller domains (residues 1-604). Neither the N-nor C-terminal β’-COP propeller domains alone were sufficient to measure binding using calorimetry (K_D_ < ~300 µM). However, when both propeller domains were present, β’-COP bound Glo3 with a K_D_ ~ 1 µM and 1:1 stoichiometry. These data suggest the Glo3 interaction requires the surface of both propeller domains, in contrast to dilysine cargo motifs, which interact only with the N-terminal propeller domain^26,27^.

### Point mutations in β’-COP or Glo3 abrogate binding *in vitro*

#### Mutagenesis mapping

We took a systematic mutagenesis approach to identify β’-COP and Glo3 residues that mediate the interaction. Sequence alignments revealed the Glo3 BoCCS region contains two highly conserved clusters of lysine residues (Figure S4A). We first tested the salt dependence of the β’-COP/Glo3 interaction by calorimetry and found high salt concentrations (500 mM NaCl) substantially weakened the interaction (Figure S4B). We concluded the interaction was likely to be mediated by electrostatic contacts. The first cluster (K233/K234/K235; Figure S4A) was proposed to interact with COPI^32^ in cell lysates, although the COPI subunit had never been identified. The second cluster (K251/K252/K255) has not been previously implicated. We tested both clusters of lysine residues (Figure 3A). The Glo3 K233E single point mutant binds β’-COP ~40-fold more weakly than does wild-type, and the Glo3 BoCCS K233E/K234E/K235E triple mutant reduces binding below detectable levels (Figure 3A). Mutating the second cluster of residues (Glo3 BoCCS K251E/K252E/K255E) also reduced binding affinity by ~40-fold. Together, these data suggest <u>both</u> basic clusters within the Glo3 BoCCS region mediate the interaction with β’-COP, and the first set of lysine residues are particularly important.

**Figure 3.**
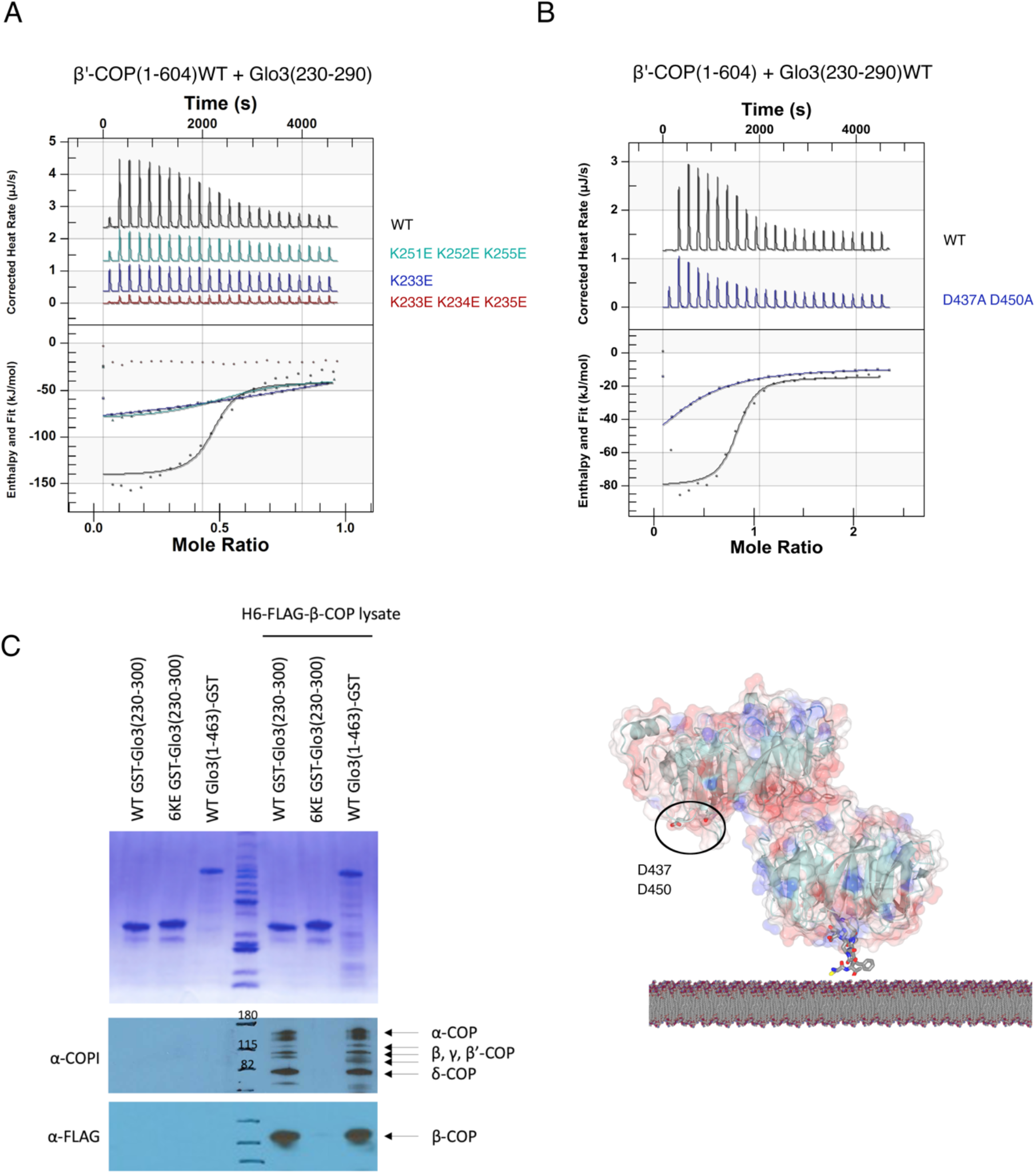
Key Glo3 lysine residues mediate an electrostatic interaction with an acidic patch on β’-COP. (A) Representative ITC experiments between wild-type β’-COP 1-604 and mutant versions of Glo3 (residues 230-290). Two Glo3 mutants substantially reduce binding in the calorimeter: a single point mutation at K233E and the K251E/K252E/K255E triple mutant. Both mutants exhibit 40-fold weaker binding as compared to wild-type Glo3 (K_D_ ~ 12 µM; Table S2). The Glo3 K233E/K234E/K235E triple mutant exhibits no measurable binding by calorimetry (K_D_ <300 µM). (B) Representative ITC experiment with wild-type untagged Glo3 (residues 230-290) and mutant β’-COP (D437A/D450A) proteins; the calculated K_D_ for this interaction is 18 µM. The D437/D450 mutant exhibits 60-fold weaker binding to Glo3, suggesting this acidic patch plays a critical role in the interaction. (C) Glo3 associates with COPI primarily using its interaction with the β’-COP subunit. Wild-type GST-Glo3 BoCCS pulled down COPI (B-and F-subcomplexes), while disrupting all six lysine residues disrupted Glo3 association with COPI. (D) β’-COP propeller domains with the acidic patch (D437/D450) circled in black; mutating this patch decreases the affinity of β’-COP for Glo3. β’-COP is shown in a membrane-bound side view with a dilysine cargo motif as grey cylinders.

Having identified key Glo3 lysine residues, we set out to find their counterparts on β’-COP, which were most likely to be acidic residues. We systematically mutated acidic patches located throughout both the N-and C-terminal propeller domains to change aspartate and glutamate residues to alanines. The vast majority of the mutant β’-COP proteins bound the Glo3 fragment with similar affinity as wild-type β’-COP (Table S2). We identified one patch on the C-terminal propeller (D437/D450) that reduced binding to Glo3 by ~60-fold when mutated to alanines (D437A/D450A; K_D_= 18 µM; Figure 3B; Figure 3D; Table S2). Several N-terminal mutant propeller proteins exhibited slightly weaker affinities (~8-fold weaker; Table S2). The N-terminal mutant with the largest effect was the R13A/K15A/R59A mutant that disrupts binding to dilysine motifs; this mutant exhibits ~16-fold weaker binding to Glo3 (Table S2; discussed further below).

Finally, we tested whether these Glo3 lysine residues are important for binding COPI in yeast (Figure 3C). Purified GST-tagged Glo3 fragments (wild-type and lysine mutants) were used as bait in pulldown experiments from budding yeast cell lysates in which both His and FLAG tags were integrated into the β-COP gene under its endogenous promoter. Full-length purified Glo3 or the Glo3 BoCCS region alone immunoprecipitated COPI from yeast cells (Figure 3C). Mutating all six lysine residues (K233E/K234E/K235E/K251E/K252E/K255E) in the Glo3 BoCCS substantially reduced the ability of Glo3 to pull down COPI. These results suggest Glo3 engages the COPI coat mainly using the β’-COP subunit. Glo3 has been proposed to interact with γ-COP appendage^41^, but these data suggest the β’-COP/Glo3 BOCCS interaction may be the primary binding site required for stable association between Glo3 and COPI.

#### Structural & modelling approaches

Unfortunately, attempts to determine a wide variety of β’-COP/Glo3 X-ray structures failed. ITC experiments revealed Glo3 binds robustly to β’-COP, so we initially used the Glo3 fragment (residues 230-290) exhibiting the greatest binding affinity for co-crystallization experiments with β’-COP. We repeatedly found the β’-COP double propeller packing within the crystal lattice excluded Glo3 fragments, which are likely to be flexible. We then systematically set up crystallization trials with Glo3 fragments and each β’-COP propeller individually; with both propellers together; with multiple synthesized Glo3 peptides of varying lengths (8-12 residues centered around two basic clusters); and using different β’-COP/Glo3 fusion constructs. We grew crystals of several protein/peptide complexes, but in each case, structure determination revealed the Glo3 peptide partially occupied the well-established dilysine cargo binding site^26,27^ (discussed further below). Attempts to grow crystals in the presence of both dilysine and Glo3 peptides were unsuccessful.

We also attempted to generate structural models of the β’-COP/Glo3 interaction using AlphaFold2 (details in Methods). Briefly, we undertook multiple runs using the β’-COP N-terminal, C-terminal, or both propellers with a variety of Glo3 fragments (summarized in Table S3; representative models in Figure S3D, S3E, S3F). AlphaFold did not converge on a solution for any attempted run. In some runs, AlphaFold placed specific Glo3 lysine residues (K234, K252, or K255) near the established β’-COP dilysine binding patch (D98/D117; Table S3). In parallel, we tested whether AlphaFold predicts the interaction between the β’-COP N-terminal propeller and dilysine motifs (Figure S3D); this served as a ‘positive control’ for computational experiments because the interaction has been observed in multiple X-ray and cryo-ET structures from different labs^26,27^. AlphaFold also consistently fails to predict the β’-COP/dilysine motif interaction. We note AlphaFold successfully predicts interactions between folded domains and short peptides in other trafficking proteins exhibiting similar affinities. One example is the SNX27 FERM interaction with DxF motifs^46,47^; AlphaFold models and the experimentally determined X-ray structure are very similar^48^. Overall, the systematic mutagenesis approach combined with biophysical experiments provided the most compelling evidence for identifying where Glo3 binds β’-COP on the C-terminal propeller (Figure 3D).

#### β’-COP binds Glo3 and dilysine motifs simultaneously *in vitro*

The biochemical and biophysical data presented here show a new direct interaction between the β’-COP subunit and the ArfGAP, Glo3. COPI sorts many important transmembrane proteins from the Golgi and endosomes, including cargoes with short amino acid motifs^49–51^ and SNARE proteins with more complex molecular signatures^6^. The β’-COP subunit recognizes dilysine motifs (KKxx or KxKxx) found in transmembrane proteins requiring recycling to the ER^26,27^. X-ray data (not shown) and some AlphaFold modeling placed Glo3 lysine residues near the dilysine binding site. We thus tested *in vitro* whether β’-COP binds dilysine cargo and Glo3 simultaneously. Both crystal structures and cryo-ET reconstructions reveal β’-COP can exist in multiple conformations, so we wondered whether Glo3 might “lock” β’-COP into a cargo-binding conformation. Alternatively, Glo3 and dilysine cargo could compete for binding to the same patch on β’-COP. However, calorimetry data (Figure S5A) showed no difference in the affinity of recombinant purified β’-COP for dilysine motifs whether Glo3 is absent or present (Figure S5A). The published β’-COP RKR dilysine binding mutant^26^ binds Glo3 with slightly weaker affinity, while the D98A/D117A mutant binds as well as wild-type protein (Figure S5B). Together, these data suggest two things. First, both Glo3 and dilysine cargo can bind β’-COP simultaneously, at least in the context of this *in vitro* assay. Second, the Glo3 binding site on the N-terminal propeller may be located near the dilysine site. The R13/K15/R59 N-terminal patch may contribute directly to binding Glo3, or alternatively, mutating it may have altered the overall electrostatic charge distribution across the propeller surface. We predict there must be a second patch on the N-terminal propeller, since both β’-COP propellers are required for binding (Figure 2B; Discussion). We could not conclusively identify the patch but current data suggest it may be located relatively close to the dilysine binding site.

#### COPI cargo sorting in yeast

We then turned to budding yeast to test whether abrogating the β’-COP/Glo3 interaction would affect Glo3 function or COPI-dependent cargo sorting *in vivo*. We tested multiple known COPI-and Glo3-dependent cargoes, including Ste2, Rer1, Emp47, and SNARE proteins.

#### Growth assays

Using the *glo3 K233E/K234E/K235E* and *glo3 K251E/K252E/K255E* mutations, we first tested if disrupting the β’-COP/Glo3 interaction would affect cell growth in a *glo3*Δ*gcs1*Δ strain background (Figure S6). We observed no difference in growth between the *glo3 K233E/K234E/K235E* and *glo3 K251E/K252E/K255E* mutants compared to the *GLO3* strain. Thus, mutations disrupting the β’-COP/Glo3 interaction do not abrogate the essential growth function of Glo3 at 30°C.

#### Ste2

Yeast α-factor receptor (Ste2) is a G-protein coupled receptor that cycles between the cell surface and internal compartments. Previous work demonstrated how post-endocytic Ste2-GFP trafficking back to the cell surface depends on the presence of Glo3^52^. We examined Ste2-GFP localization in *glo3*Δ*gcs1*Δ cells expressing wild-type *GLO3, glo3 K233E/K234E/K235E* or *glo3 K251E/K252E/K255E* mutants (Figure 4A). We observed a significant difference in the amount of Ste2-GFP at the plasma membrane in *GLO3* versus *glo3* mutant cells (Figure 4B). Less Ste2-GFP was observed at the plasma membrane in both *glo3* mutant strains, and mis-localization to the vacuole was apparent (Figure 4A).

**Figure 4.**
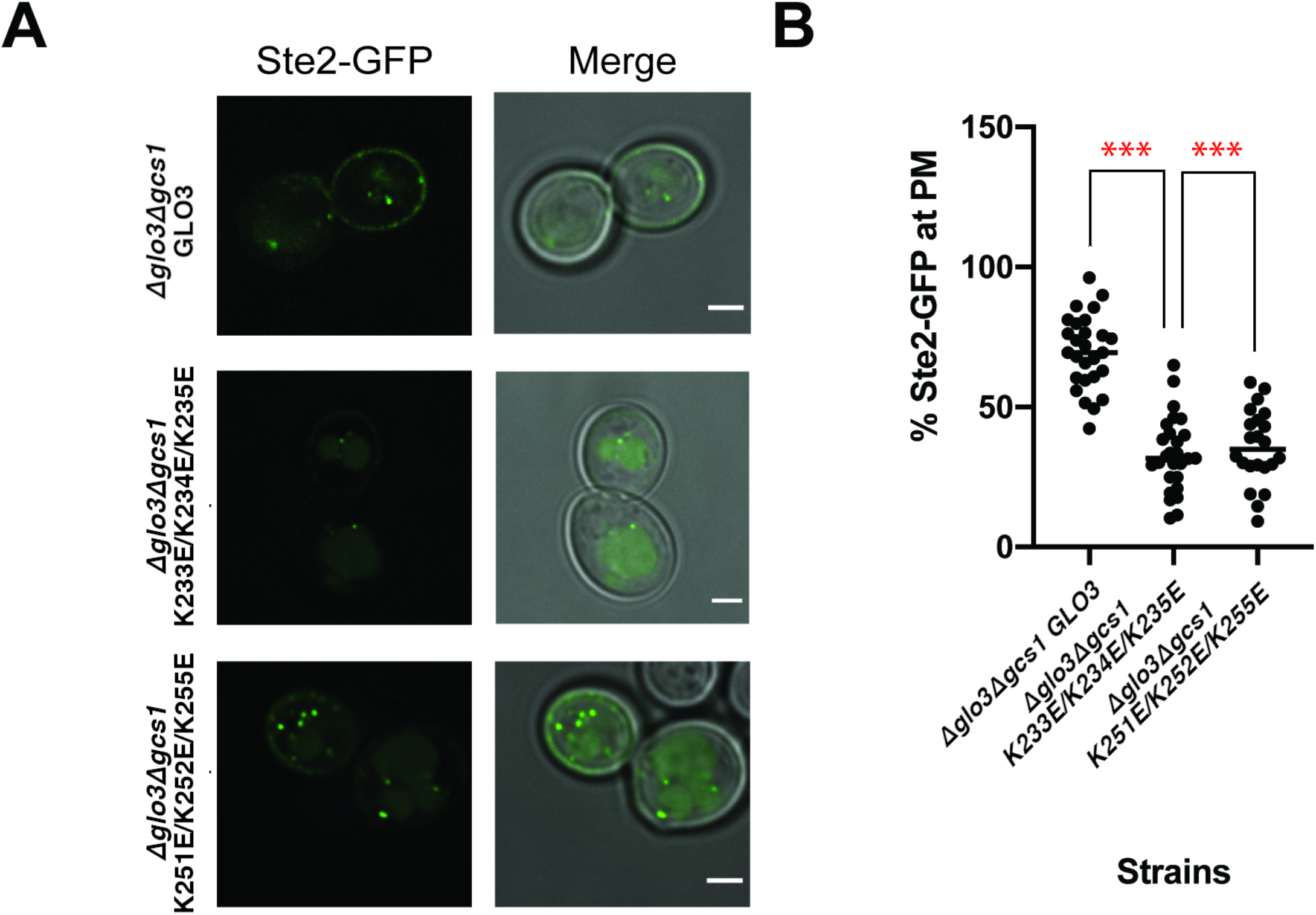
Ste2 is mis-sorted to the vacuole when the β’-COP/Glo3 interaction is disrupted in *S. cerevisiae*. (A) Fluorescence imaging of Ste2-GFP in *glo3ΔΔgcs1ΔΔ* strains with *GLO3* or *glo3* mutants (K233E/K234E/K235E or K251E/K252E/K255E) introduced on a plasmid. Scale bar represents 2 µm. (B) Box plots showing percentage of Ste2-GFP observed at the plasma membrane in each strain with median marked (black line). Mutating either lysine cluster in yeast cells causes a significant difference in Ste2-GFP sorting compared to the *GLO3* strain. Statistical comparisons were pairwise between *GLO3* and mutants; data were analyzed using a one-way ANOVA (Prism); and probability values of less than 0.001 are represented by ***.

#### Emp47

α-and β’-COP recognize dilysine sorting motifs (KKxx or KxKxx) present at the C-terminus of cargo proteins. Emp47 is an established COPI cargo containing a KxKxx motif that can be recognized by β’-COP; it is therefore a suitable cargo to study COPI-mediated Golgi-to-ER retrieval involving the β’-COP subunit. Mutations that disrupt Emp47 sorting into COPI vesicles cause Emp47 mis-localization to the vacuole where it is degraded. We examined the stability of Emp47-myc over time in *Δglo3Δgcs1* cells after inhibiting new protein synthesis with cycloheximide, but we did not observe an effect on Emp47 sorting. Emp47 appeared to remain stable over time in *glo3 K233E/K234E/K235E* and *glo3 K251E/K252E/K255E* cells (Figure S7A).

#### Rer1

We next tested whether disrupting the β’-COP/Glo3 interaction would affect two different protein cargoes known to cycle between the Golgi and ER in COPI coated vesicles. We first examined localization of GFP-Rer1 (Figure S7B), a protein that localizes to early Golgi cisternae at steady-state but rapidly cycles between the ER and early Golgi in COPI and COPII vesicles. Disruption of COPI function causes mis-localization of Rer1-GFP to the vacuole^53^. However, we found no difference in Rer1-GFP localization between the *glo3* mutants and wild-type *GLO3* in the strains tested. Together, data on Rer1 and Emp47 suggest the *glo3* basic patch mutants do not appear to disrupt COPI function in retrieving cargoes from the Golgi to ER.

#### SNAREs

The yeast R-SNARE, Snc1, is an exocytic SNARE that cycles between the Golgi and plasma membrane and has been shown to be mis-localized when Glo3 is deleted^52,54^. We examined mNG-Snc1 localization in *glo3 K233E/K234E/K235E* and *glo3 K251E/K252E/K255E* mutants in the *glo3*Δ*gcs1*Δ background (Figure S8A/B/C). mNG-Snc1 normally localizes to punctate structures inside the cell, as well as to the plasma membrane. We found no significant difference in the amount of mNG-Snc1 at the plasma membrane relative to internal structures. However, in the *glo3 K233E/K234E/K235E* cells, mNG-Snc1 localizes to ring or tubular structures that are less frequently observed in cells expressing wild-type *GLO3* (Figure S8A). These structures may indicate abnormal Golgi and/or endosome morphology. This significant phenotype suggests disrupting the β’-COP/Glo3 interaction affects COPI function required to maintain Golgi morphology.

Bet1 is a Q-SNARE involved in ER-Golgi trafficking that localizes to early Golgi compartments at steady-state and also interacts with the BoCCS region of Glo3^32,55^. We examined mNG-Bet1 localization (Figure S8D/E/F) in *glo3*Δ*gcs1*Δ cells expressing wild-type *GLO3* or the basic cluster mutants. We identified a small number of abnormal ring structures in *glo3 K233E/K234E/K235E* cells. Data from the mutants were not significantly different from *GLO3* cells (Figure S7E/F); the number of abnormal structures was on the edge of significance, in contrast to data on mNG-Snc1.

Together, data from multiple COPI transmembrane receptor cargoes suggest the β’-COP/Glo3 interaction is more important for cargoes recycling via the endosome/plasma membrane and less important for cargoes cycling between the *cis*-Golgi and ER (see Discussion).

## Discussion

### Molecular details of the β’-COP/Glo3 interaction

Published data^36^ previously demonstrated how Glo3, but not Gcs1, stably associates with COPI using affinity purification from yeast cell lysates. Here we show for the first time that Glo3 and the β’-COP subunit exhibit a direct molecular interaction with a low micromolar K_D_. We mapped specific residues in both Glo3 and β’-COP propeller domains to identify an acidic patch (D437/D450) on the β’-COP C-terminal propeller that binds residues located in the Glo3 BoCCS region. The Glo3 BoCCS contains two basic lysine clusters that interact directly with β’-COP (Figure 3A). These two clusters are separated by ~20 amino acids and could span up to 50 Å if this region is unconstrained by secondary structure. We have shown both β’-COP propeller domains are required to bind Glo3 BoCCS (Figure 2B). Taken together, it is tempting to speculate each Glo3 cluster binds an acidic patch on each β’-COP propeller domain. This implies there is a second acidic patch located on the N-terminal propeller. We were unable to identify this patch using a systematic mutagenesis approach, but circumstantial evidence implies a patch may be located near the R15/K17/R59 patch (Figure S5B). Attempts to determine structures experimentally or to use AlphaFold to generate structural models for the β’-COP/Glo3 interaction failed. Crystallography attempts were confounded by two primary factors. First, the minimal Glo3 fragment required to maintain the binding affinity is relatively long (50-60 amino acids), which makes it a difficult co-crystallization target. Second, short Glo3 peptides used in co-crystallization experiments are similar to dilysine binding motifs in cargo molecules that bind the N-terminal β’-COP propeller. These peptides partially occupied the established dilysine binding site (data not shown). *In vitro* calorimetry data (Figure S5) suggests Glo3 and dilysine cargo occupy distinct binding sites on the β’-COP propellers, so the structural data are very likely to be crystallization artefacts. It is more challenging to speculate why AlphaFold did not yield testable models. One possibility is lack of sequence conservation among ArfGAPs. ArfGAP flexible linkers vary in both sequence and length among species, and there are multiple basic residues in the linkers. Algorithms may struggle to predict which clusters are conserved among species. Furthermore, it has not yet been shown that mammalian ArfGAP2/3 interacts with β’-COP in the same way, so it remains possible that mammalian COPI assembly differs from yeast. New structural approaches will be required to uncover precise molecular details of the β’-COP/Glo3 interaction, particularly to identify residues on the N-terminal propeller.

### β’-COP emerges as a molecular platform

Multiple lines of evidence now suggest the β’-COP subunit acts as a molecular platform to coordinate binding to multiple protein partners within the COPI coat. Propeller domains in other trafficking proteins also act as platforms. For example, clathrin contains a single propeller called terminal domain (TD), which contains four binding sites for binding partners^42,44,56,57^. The COPI coat contains four propeller domains: two in β’-COP and two in α-COP. Until now, only one specific binding site on both α-and β’-COP propeller domains has been identified: the N-terminal propeller domains bind short dilysine motifs (KKxx or KxKxx) found in transmembrane protein cargoes cycling between the Golgi and ER^22,26,27^. β’-COP is located near γ-appendage in the F-subcomplex^21^ in reconstituted coats, and β’-COP binds ubiquitinated Snc1^6^, although high-resolution structural data are not available. β’-COP thus directly engages multiple important proteins in assembling COPI coats.

In this work, we identify two new direct binding partners for β’-COP: the Glo3 BoCCS region and Arf1 (Figure 1). We combine data on this new β’-COP/Glo3 BoCCS interaction with a published Glo3 GAP/Arf1 structural model^58^ to propose how β’-COP may bind Glo3 and Arf1 on membranes in the presence of cargo (Figure 5). Multiple lines of evidence support this model. In cryo-ET reconstructions, β’-COP is located adjacent to both Arf1^20^ and the ArfGAP2/3 GAP domain^21^, although these structures did not report molecular details of a β’-COP/Arf1 interface. Pulldown data from this work demonstrate biochemically how β’-COP propeller domains can directly bind Arf1 and Glo3, and we ascribe the first function to the C-terminal β’-COP propeller domain. One caveat to this model is that the orientation of the BoCCS region remains unknown. The model presented here is drawn based on reported interactions, including an interaction with-COP appendage^41^ in mammalian cells. Together, data suggest β’-COP may coordinate key events in COPI coat function, perhaps coupling cargo recognition with F-subcomplex, Arf1, and ArfGAP binding. It will be critical to test in future experiments whether cargo binding by β’-COP influences Arf1(GTP) hydrolysis by Glo3.

**Figure 5.**
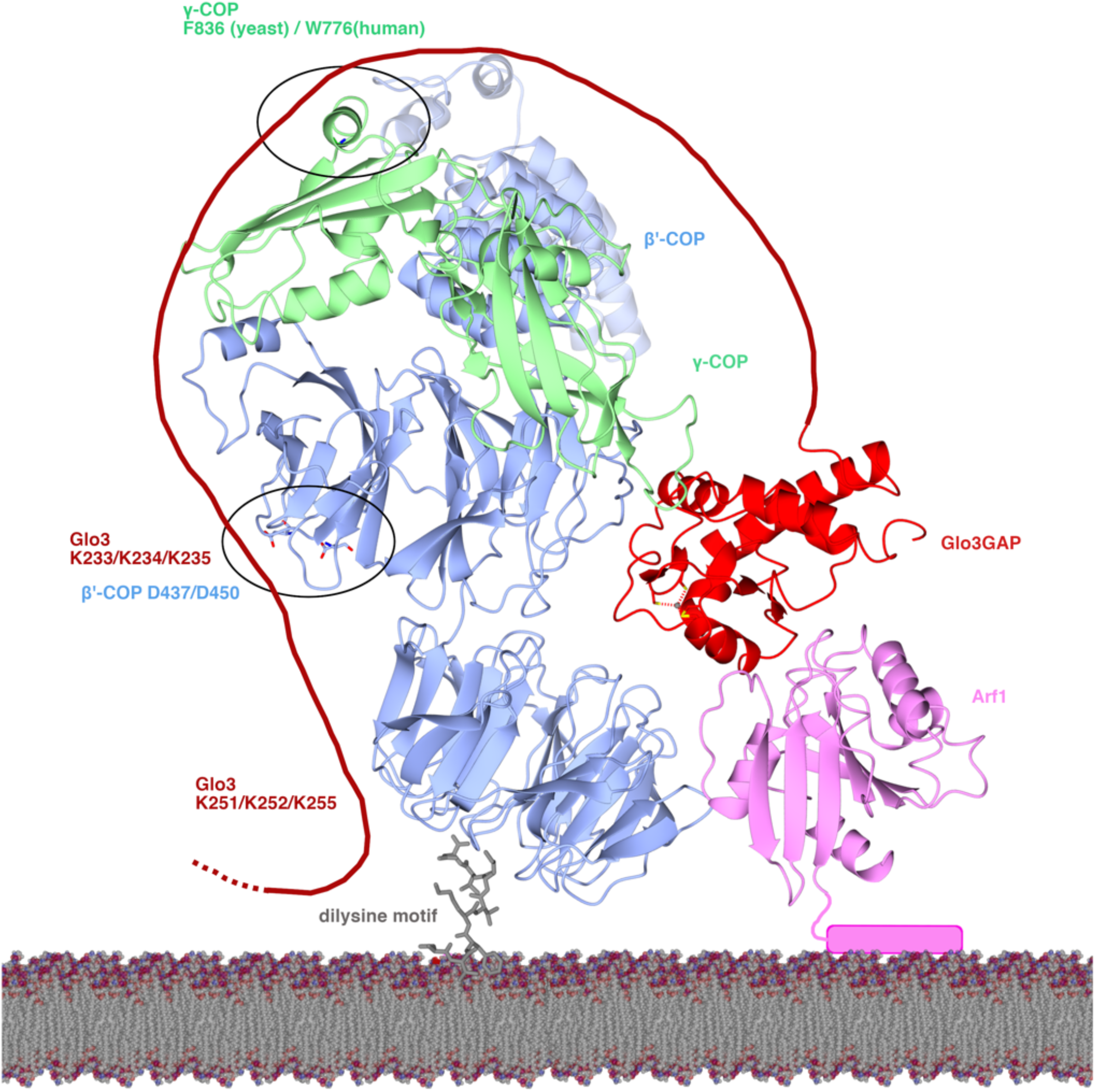
γ’-COP is a molecular platform on Golgi membranes. Model for the interaction between γ’-COP (blue ribbons; WD-repeat domains and solenoid shown); Arf1 (pink ribbons with N-terminal amphipathic helix shown as cylinder); γ-COP appendage (green ribbons); and Glo3 (GAP domain as red ribbons and BoCCS region as red line). The dashed line marks the start of Glo3 GRM region; its position and orientation remain unknown, but it is predicted to have a C-terminal amphipathic helix that may insert into the membrane. The dilysine motif in transmembrane cargoes is shown as grey cylinders. The position of γ-COP was generated based on cryo-ET reconstructions of COPI (PDB: 5NZS). The first Glo3 lysine cluster (K233/K234/K235) may interact with the D437/D450 patch on the C-terminal β’-COP propeller, and the second cluster (K251/K252/K255) may interact with the N-terminal β’-COP propeller (discussed further in text). γ-COP appendage also binds Glo3; residue F836 is implicated in binding, but the Glo3 residues remain unknown.

### Cargo sorting and Golgi morphology

ArfGAP proteins have long been known to play important roles in membrane trafficking. Yeast tolerate the loss of either Gcs1 or Glo3, but not both proteins. Despite their importance, the molecular roles of ArfGAP proteins have been difficult to define. This study defines a new molecular interaction between β’-COP and Glo3, which can now be used to generate molecular tools for separation of function studies to probe COPI function with Glo3 and Gcs1 more precisely.

When the β’-COP/Glo3 interaction was disrupted in budding yeast, we observed two distinct phenotypes. First, Ste2 (α-factor receptor) exhibited a trafficking defect at steady state and tended to be sorted to the vacuole instead of cycling via the plasma membrane. Second, we observed defects in Golgi morphology. Published data have implicated Glo3 in Snc1 localization^52^ because loss of the entire BoCCS region partially traps Snc1 in internal punctate structures. Here, we imaged tagged SNARE proteins (Snc1 or Bet1) and found the Golgi sometimes formed abnormal structures (Figure S7) when we specifically disrupted the β’-COP/Glo3 interaction. In contrast, we did not identify mis-trafficking of well-characterized COPI cargoes that cycle between the *cis*-Golgi and ER, such as Rer1 and Emp47. β’-COP can bind both Glo3 BoCCS region and dilysine motifs simultaneously *in vitro* (Figure S5), which supports the idea that β’-COP may directly engage Glo3 and some cargoes simultaneously. However, binding Glo3 does not appear to be required for proper dilysine cargo sorting in the *cis*-Golgi (Figure S6). Together, these data may implicate the β’-COP/Glo3 interaction in maintaining proper steady-state levels of receptor cargoes (e.g. Ste2) and SNAREs (e.g. Snc1) that cycle from endosomes via the TGN to the plasma membrane. These data may also suggest a separation of function between Glo3 and Gcs1, which will be important to test further.

### Implications and future directions

Our data raise multiple new questions about how and why the COPI coat couples Glo3 binding to Arf1 hydrolysis and recognition of cargo and SNARE proteins. Classically, Arf1(GTP) has been reported to recruit the F-subcomplex^17,19,20^. Goldberg and colleagues showed how the β/δ-COP and γ/ζ-COP sub-complexes interact directly with Arf1(GTP) to promote an “open” conformation analogous to AP-1/AP-2^59,60^ in clathrin coated vesicles. Our data suggest β’-COP shows no preference for either Arf1 nucleotide-bound state *in vitro*, so it is possible only the F-subcomplex “senses” Arf1 state to promote the open or hyper-open conformations observed in crystal structures^19^ and reconstituted vesicles^20,21^. We speculate the relatively weak β’-COP/Arf1 interaction could further stabilize an assembling coat, and the interface must be located on the opposite face to the Arf1 switch I/II regions. The tripartite interaction between β’-COP, Glo3, and Arf1 bears further investigation to understand how Arf1 hydrolysis mediated by Glo3 may be coupled to cargo and/or SNARE binding.

Multiple lines of evidence suggest the Glo3 BoCCS region plays an important regulatory role in COPI trafficking. The BoCCS region has been shown to be sufficient for rescuing growth defects in yeast strains lacking both *GLO3* and *GCS1*^54^. Arakel and colleagues^36^ demonstrated re-introducing the BoCCS region alone rescues growth in the Glo3 GAP-dead mutant (R59K) strain. The BoCCS region co-immunoprecipitates Emp47, Bet1, and Bos1^54^. This may represent a direct interaction, but our data suggest it may alternatively reflect the ability of Glo3 BoCCS to interact with Emp47^26,27^ indirectly via β’-COP. In addition, Glo3 has been implicated in Sec22 binding^61^ but the molecular details remain unknown. Based on both published and new data reported here, we speculate the β’-COP/Glo3 interaction may be necessary to ensure proper incorporation and recycling of specific receptor and SNARE proteins, even in the absence of a functional Glo3 GAP domain.

## Acknowledgements

We sincerely thank David Owen and members of the Jackson and Graham labs for helpful feedback, discussion, and critical reading of the manuscript. BX, CJ, and AK performed protein expression, protein purification, biochemical, and biophysical experiments. BX, CG, SD, and JB undertook yeast experiments, including growth and cargo sorting assays, and imaging experiments. BX and LPJ wrote the paper with input from all authors. LPJ conceived the project. BX, CJ, AK, and LPJ are supported by NIH R35GM119525. LPJ is a Pew Scholar in the Biomedical Sciences, supported by the Pew Charitable Trusts. CG, JB, SD, and TRG are supported by NIH R01GM118452. The authors declare no competing conflicts of interest.

## Materials and Methods

### Antibodies

The following antibodies were used in this study: mouse α-myc (Invitrogen, 9E10); rabbit α-Arf1 (from Todd Graham, Vanderbilt University); α-His (Abcam, NB100-63173), mouse α-CPY (Invitrogen 10A5B5).

### Cloning and mutagenesis

For structural and biochemical analyses, C-terminal His-tagged β’-COP and Glo3 were placed into the Nde I/ BamHI or Nde I /HinD III sites of in-house vector pMW172^62^ under control of the T7 promoter. N-terminal His-tagged yeast Arf1 was placed into pET-21a (+) vector in the Nde I/ BamHI sites. The Glo3 BoCCS and GRM regions and β’-COP (residues 300-604) were ligated into BamHI/ NotI sites of pGEX-6P-1 (GE Healthcare), resulting in an N-terminal GST-tagged proteins with a 3C-protease cleavage site. Other GST-tagged Glo3 and β’-COP constructs were sub-cloned into Nde I/ HinD III or Nde I/ BamHI sites of pMWGST, a modified form of pMW172 incorporating a C-terminal, thrombin cleavable GST tag. A two-stage quick-change mutagenesis protocol was used to introduce mutations in Glo3 and β’-COP. In this protocol, mutagenic primers (Sigma) were created for the desired mutations. In the first step, two polymerase chain reactions (PCRs), with either the mutagenic 5’ or 3’ primer, were amplified around the plasmid. The two reactions were then combined in an additional PCR step, followed by Dpn I digest and transformation. All constructs were verified by sequencing (GENEWIZ) prior to use.

### Protein expression and purification

All constructs were expressed in BL21(DE3)pLysS cells (Invitrogen) for 16-20 hours at 22°C following induction with 0.4 mM IPTG. His-tagged Arf1constructs were purified in 20 mM Tris-HCl (pH 8.0), 200 mM NaCl, 0.5 mM TCEP, and 5 mM MgCl_2_. Full-length Glo3 was purified in 20 mM HEPES (pH 7.5), 500 mM NaCl, and 1 mM DTT. Other Arf1 and Glo3 constructs and all β’-COP constructs were purified in 20 mM HEPES (pH 7.5), 200 mM NaCl, and 1 mM DTT. AEBSF (Calbiochem) was incorporated at early stages of all purifications. Cells were lysed by a disruptor (Constant Systems Limited) and proteins were affinity purified using glutathione sepharose (GE Healthcare) or HisPur cobalt resin (Thermo Scientific) in buffers listed above. GST-tagged proteins were cleaved overnight with thrombin (Recothrom, The Medicine Company) or 3C-protease (made in-house) at 4°C and eluted in batch. All proteins were further purified by gel filtration on a Superdex S200 preparative or analytical column (GE Healthcare).

### GST pulldown assays

GST or GST-tagged β’-COP (residues 1-604) or Glo3 constructs (GAP domain, BoCCS region, GRM region) were immobilized on glutathione sepharose resin (GE Healthcare) for one hour on ice. The resin was incubated for one hour on ice with prey proteins (Glo3-His8; T31N or Q71L Arf1His6; or β’-COP604His6) in 20mM HEPES (pH7.5), 200mM NaCl, 2mM DTT, and 0.5% NP40. Samples were washed three times with the same buffer. Proteins were eluted from the resin using buffer plus 30 mM reduced glutathione. Gel samples were prepared from the supernatant following elution, and the assay was analyzed by Commassie staining of SDS-PAGE gels. The gels were further analyzed by western blotting, using anti-His (Abcam, NB100-63173) and rabbit anti-Arf1 antibodies (from Todd Graham, Vanderbilt University).

### Isothermal titration calorimetry

ITC experiments were conducted on a NanoITC instrument (TA Instruments) at 20°C. The molar concentration of protein in the syringe was at least 5 times that of protein in the cell. All experiments were carried out in 10 mM HEPES (pH 7.5), 100 mM NaCl, and 0.5 mM TCEP, filtered and degassed. Incremental titrations were performed with baseline of 100 seconds and injection intervals of 200 seconds. Titration data was analyzed in NANOANALYZE (TA instruments) to obtain a fit and value for stoichiometry (n) and equilibration association constant (K_a_). K_D_ values were then calculated from the association constant.

### Sequence alignments

In order to map residue conservation of Glo3, sequences of full-length and BoCCS region alone of Glo3 from *S. cerevisiae, M. musculus, H. sapiens, D. melanogaster, C. elegans, S. pombe, A. thaliana* were aligned using ClustalW^63^ and Praline^64^. A partial alignment is shown in Figure S4 to highlight key conserved lysine residues in yeast, murine, and human proteins required to interact with β’-COP.

### Yeast strains and plasmids

Standard media and techniques for growing and transforming yeast were used. Glo3 mutant strains were constructed by plasmid shuffling (PXY51) on 5’-fluoro-orotic acid (5-FOA) plates. Yeast strains used in this study are listed in Table S4. Plasmids constructions were performed using standard molecular manipulation. Wild-type *GLO3* gene were cloned into pRS315 yeast vectors with endogenous promoter and terminator sequences. Mutations were introduced using the two-stage quick-change mutagenesis protocol.

### Yeast growth assays

Strains containing pRS416-*GLO3* (wild-type) and pRS315-*glo3* (mutant constructs) were grown at 30°C. Cells were then sub-cultured, and an equal OD of each strain was loaded into a 96-well plate. Using a prong replicator, strains were stamped onto both appropriate synthetic media and 5-FOA plates to select against the pRS416-*GLO3* plasmid. Cells were grown for four days and images of these plates were captured.

### Fluoresence imaging

Three biological replicates of transformed strains were sub-cultured for imaging in appropriate synthetic media. Cells at mid-log phase were then mounted on glass slides and observed immediately at room temperature. Images were acquired using a DeltaVision Elite Imaging System (GE Healthcare Life Sciences, Pittsburgh, PA) equipped with a 100x objective lens. Z-stack of images were collected for Green channel and DIC. All images were deconvolved using softWoRx software (GE Healthcare Life Sciences). Cells were chosen using green fluorescence expression.

### Data analysis and statistics

To quantify levels of Ste2-GFP at the cell surface, traces were drawn just outside and inside the plasma membranes using the freehand drawing tool in ImageJ to measure the total cellular fluorescence (Cell_fl_) and Internal fluorescence (Int_fl_), respectively. The percent of Ste2-GFP at the plasma membrane was then calculated by (Cell_fl_ - Int_fl_)/Cell_fl._ Statistical differences were determined using a one way ANOVA test in GraphPad Prism version 8.0 (GraphPad Software, San Diego, California, USA, www.graphpad.com). Images of flourescently tagged SNARE proteins were coded, and the number of tubules, puncta, and rings were counted in a blinded experiment. Statistical differences against wild-type were determined using Mann-Whitney test, and data were visualized using Python. Probability values of less than 0.04, 0.01, or 0.001 were used to show statistically significant differences and are represented with *, **, or *** respectively.

### Dilysine reporter construct trafficking in yeast

Yeast Emp47 under control of its endogenous promoter was used as the backbone for reporter constructs, and a myc-tag was incorporated following the transmembrane domain to facilitate Western blotting. The KxKxx reporter contained endogenous Emp47 transmembrane domain followed by the cytoplasmic sequence (RQEIIKTKLL). The reporter was introduced into appropriate *GLO3* wildtype and *glo3* mutant strains. Reporter levels were monitored by Western blotting against the myc tag both prior to and following a 20 μg/mL cycloheximide chase at time points equal to 0, 30, and 60 minutes. Cells were lysed using glass beads with SDS buffer and then boiled at 95°C for 5 minutes. Western blots were probed with mouse monoclonal anti-myc (Invitrogen 9E10) for the reporter and with mouse anti-CPY (Invitrogen 10A5B5) as a loading control. The membrane was then incubated with HRP-goat anti-mouse IgG secondary antibody (Invitrogen) or IRDye 800CW goat anti-mouse IgG secondary antibody (LI-COR) to visualize.

### Structural predictions using AlphaFold

To generate structural models of yeast Glo3 and β’-COP proteins, we used both AlphaFold2 Multimer neural-network ^65,66^ implemented within the freely accessible ColabFold pipeline^67^ and standalone AlphaFold2 Multimer installed through SBGrid. Sequences of single or double propeller domains of β’-COP were used in complex with different Glo3 lengths. For each structural prediction experiment, sequences from evolutionarily related protein in the form of a multiple sequence alignment were created by Jackhammer^68^ in local AlphaFold or by MMseqs2 in ColabFold^67^, and AlphaFold2 model building was executed using standard settings. Structural relaxation of final protein complexes was performed with Amber to generate five models per protein complex.

## Supplemental figures

**Figure S1.**
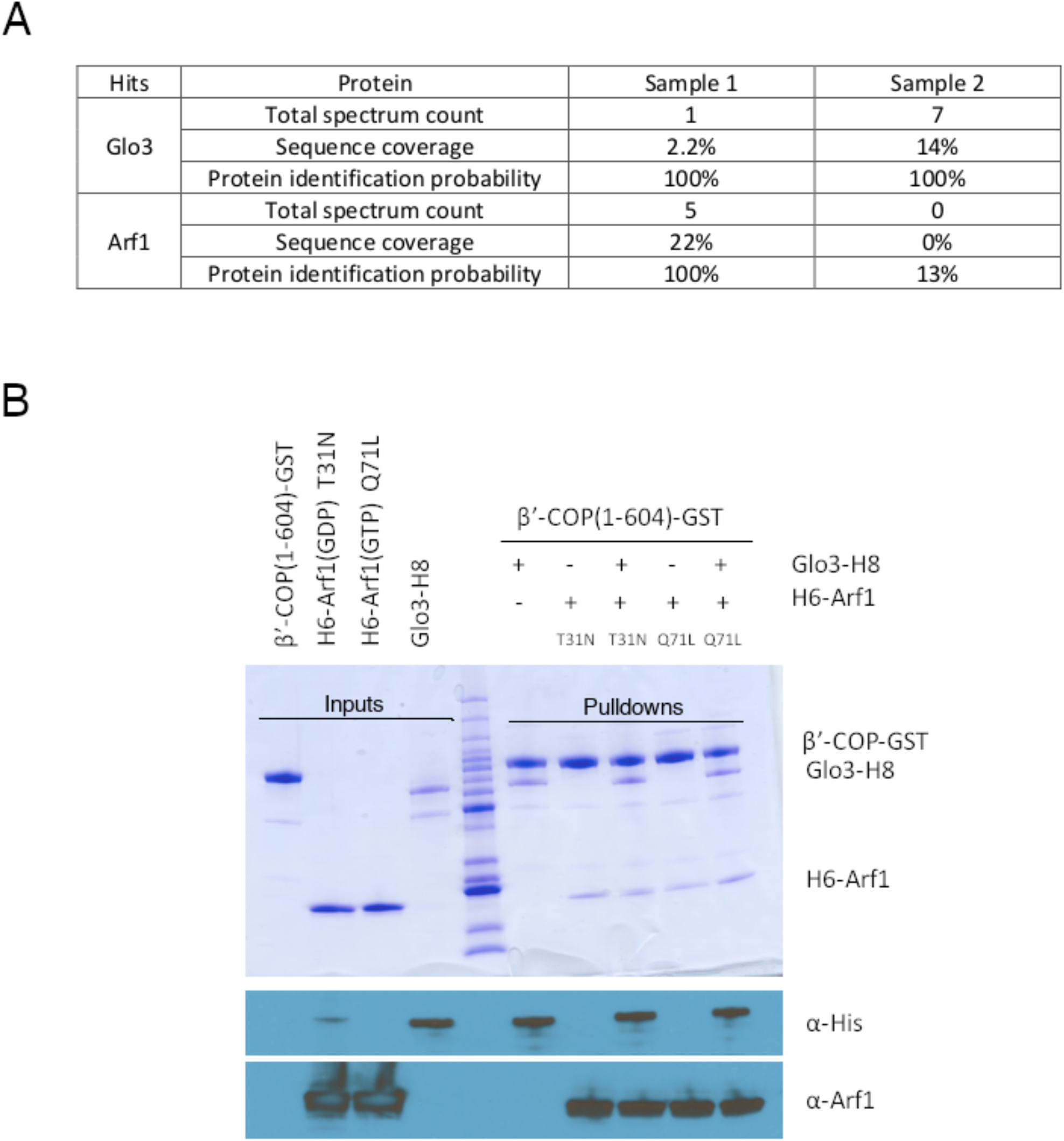
β’-COP, Arf1, and Glo3 form a ternary complex *in vitro*. (A) Glo3 and Arf1 were identified as potential direct binding partners using GST-tagged β’-COP protein as bait in yeast cell lysates. The table summarizes mass spectrometry results for these two hits. (B) GST-pulldown experiments using GST-tagged β’-COP propeller domains (residues 1-604 with C-terminal GST tag) as bait and full-length Arf1-H6 with and without full-length Glo3-H8 as prey. We tested binding to both nucleotide-bound forms of Arf1: GDP-locked (T31N) or GTP-locked (Q71L). β’-COP can pull down both Arf1 and Glo3 simultaneously. β’-COP does not appear to show a preference for Arf1 nucleotide state; it pulls down Arf1 T31N or Q71L equally well when Arf1 is added at a 5:1 molar ratio.

**Figure S2.**
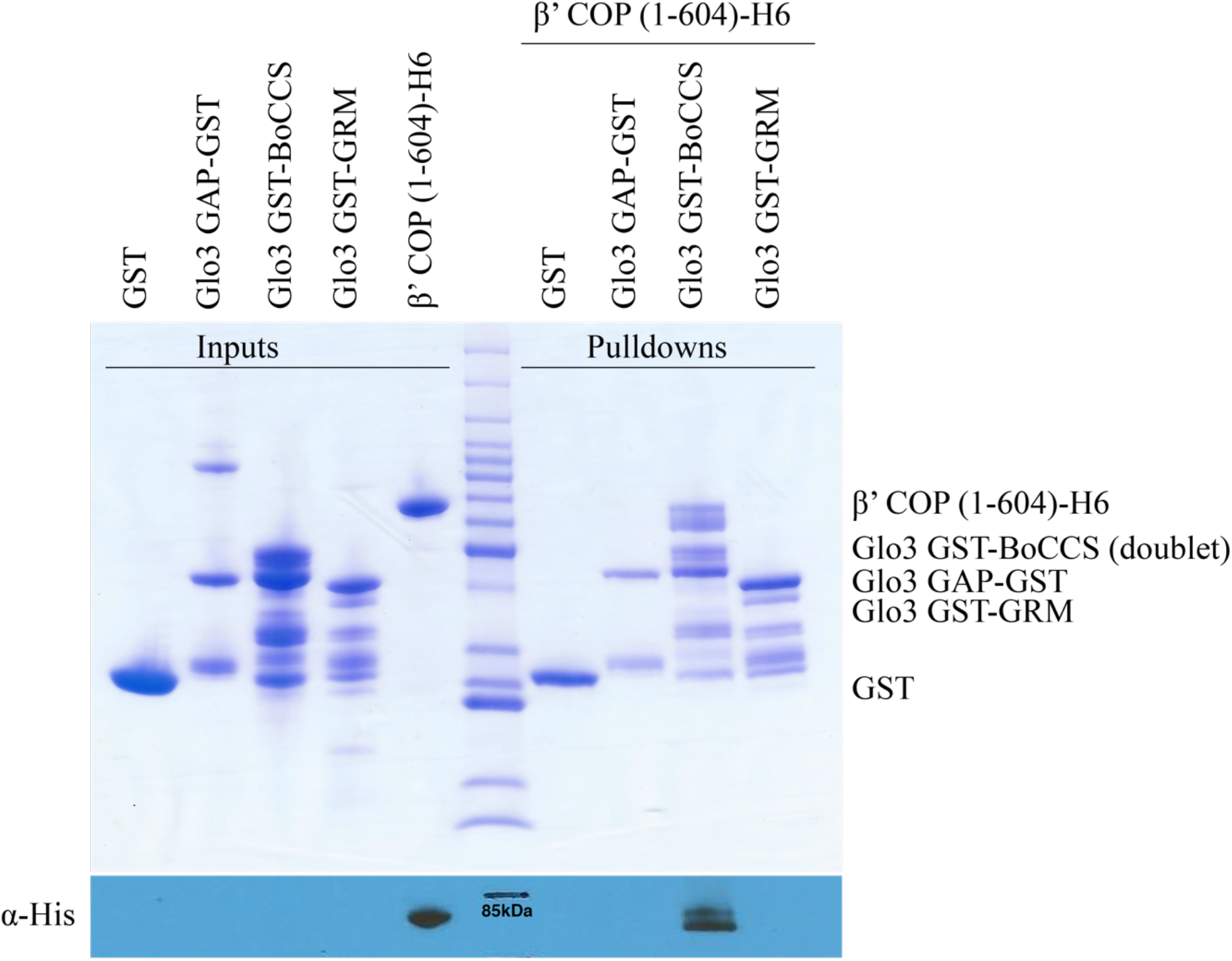
β’-COP WD-repeat domains directly bind the Glo3 BoCCS region *in vitro*. GST-tagged Glo3 fragments (GAP-GST, residues 1-150; GST-BoCCS, residues 208-383; or GST-GRM, residues 350-493) were used as bait with β’-COP-H6 to determine which portion of Glo3 binds β’-COP. Only the GST-BoCCS fragment exhibited direct binding. Both the BoCCS and GRM fragments are unstable in solution, most likely because they contain long regions predicted to be unstructured. Mass spectrometry data (not shown) confirms the top two bands in the GST-BoCCS input lane correspond to Glo3 BoCCS peptides.

**Figure S3.**
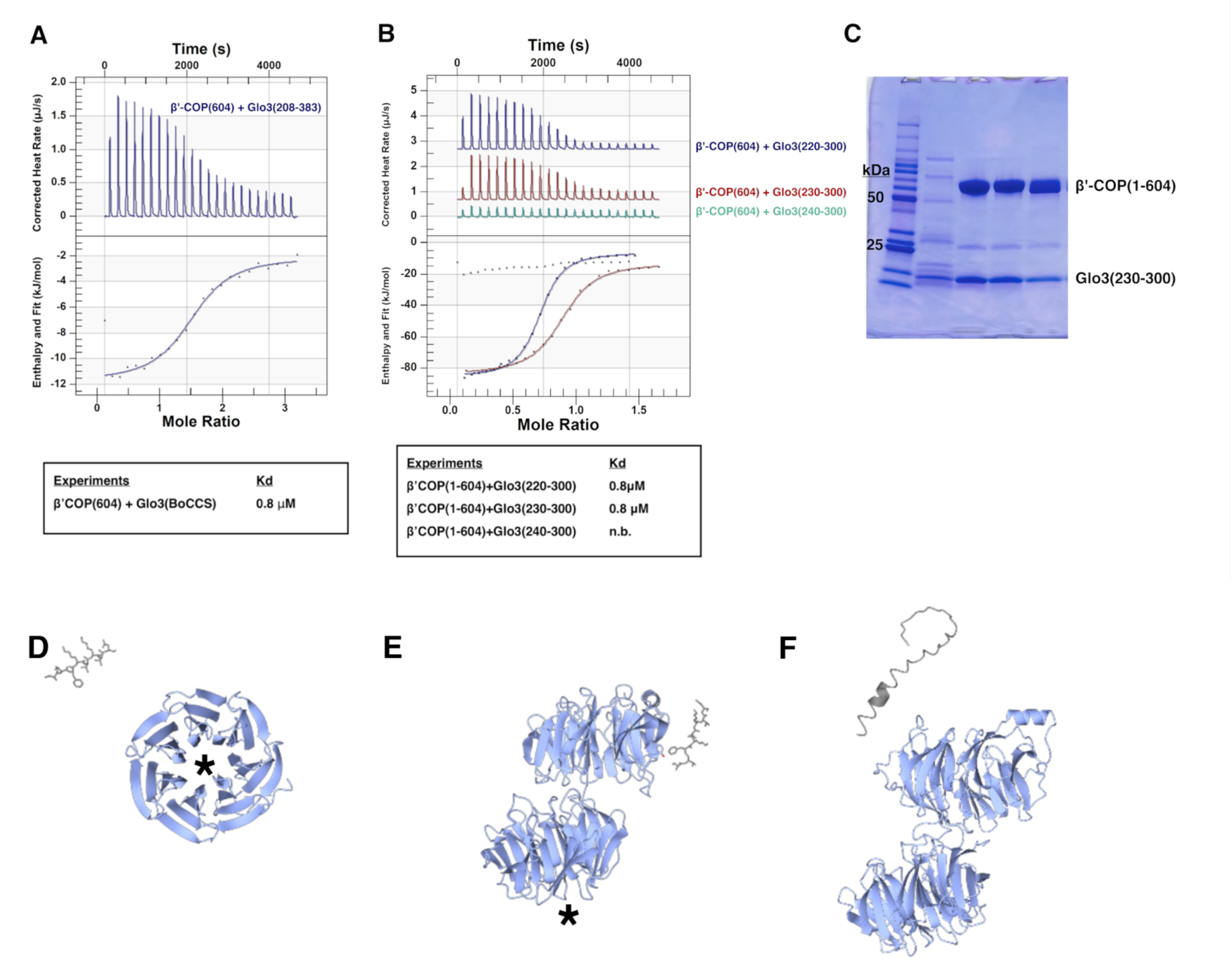
The Glo3 BoCCS region interacts directly with β’-COP. (A) An ITC experiment with full Glo3 BoCCS fragment (residues 208-383) shows a low micromolar K_D_. (B) ITC experiments to ascertain minimum Glo3 fragment required for binding. Glo3 N-terminal residues 230-240 are required for measurable binding in the calorimeter, while removing residues 220-230 exhibits no measurable effect on binding affinity. “n.b.” denotes no measurable binding in the calorimeter (K_D_ > 300µM). (C) Representative SDS-PAGE gel of β’604/Glo3 (residues 230-300) complex following purification; the complex elutes together over gel filtration columns (data not shown). Although some Glo3 fragments are unstable over time, recombinant fragments used for ITC runs exhibit high levels of purity and stability. (D-F) Representative models generated by AlphaFold2. The well-established interaction between β’-COP and dilysine motifs should serve as a ‘positive control’ computational experiment, but AlphaFold failed to predict binding between dilysine motifs (grey cylinders) and the N-terminal β’-COP propeller (D) or to both propeller domains (E) shown in blue ribbons. The known dilysine binding site is marked by black asterisks. (F) One representative model from an AlphaFold experiment with β’-COP 1-604 (blue ribbons) and Glo3 residues 230-290 (grey ribbons). Results from modelling experiments are reported in Table S3.

**Figure S4.**
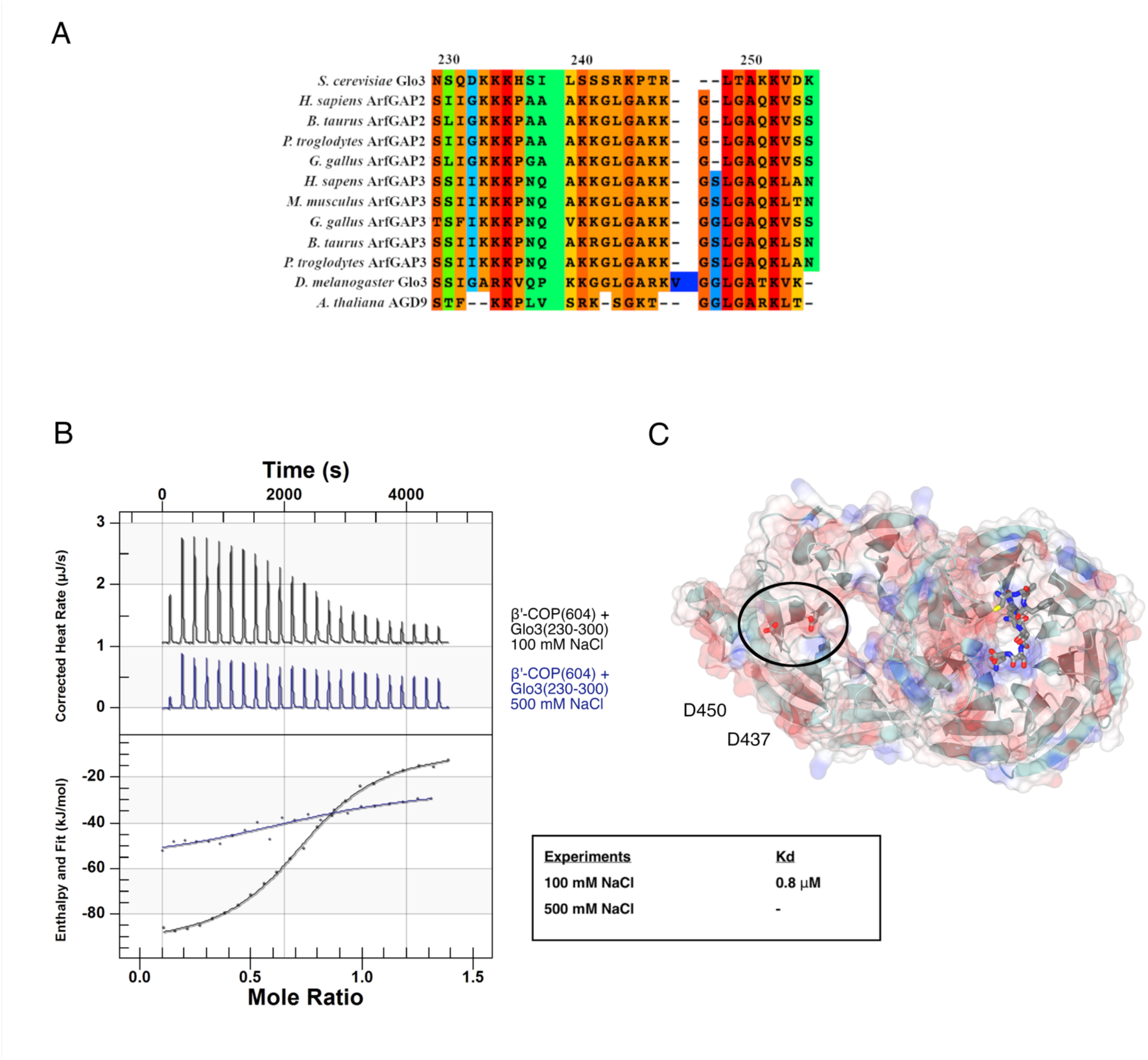
Conserved Glo3 lysine residues mediate an electrostatic interaction with β’-COP C-terminal propeller domain. (A) Glo3 partial sequence alignment highlighting key conserved residues between *S. cerevisiae* Glo3 and mammalian ArfGAP2/3 homologs in *M. musculus* and *H. sapiens*. Glo3 residues 230-300 are labeled. (B) ITC experiment between wild-type β’-COP residues 1-604 and Glo3 residues 230-270 in 100 mM NaCl and 500 mM NaCl. Near physiological salt concentrations, the two proteins interact with a low micromolar K_D_. The same experiment in high-salt buffer (500 mM NaCl) disrupts the interaction, suggesting electrostatic residues play an important role. In high salt, binding was too weak to determine a K_D_. (C) View up from membrane showing the β’-COP D437/D450 acidic patch that binds Glo3.

**Figure S5.**
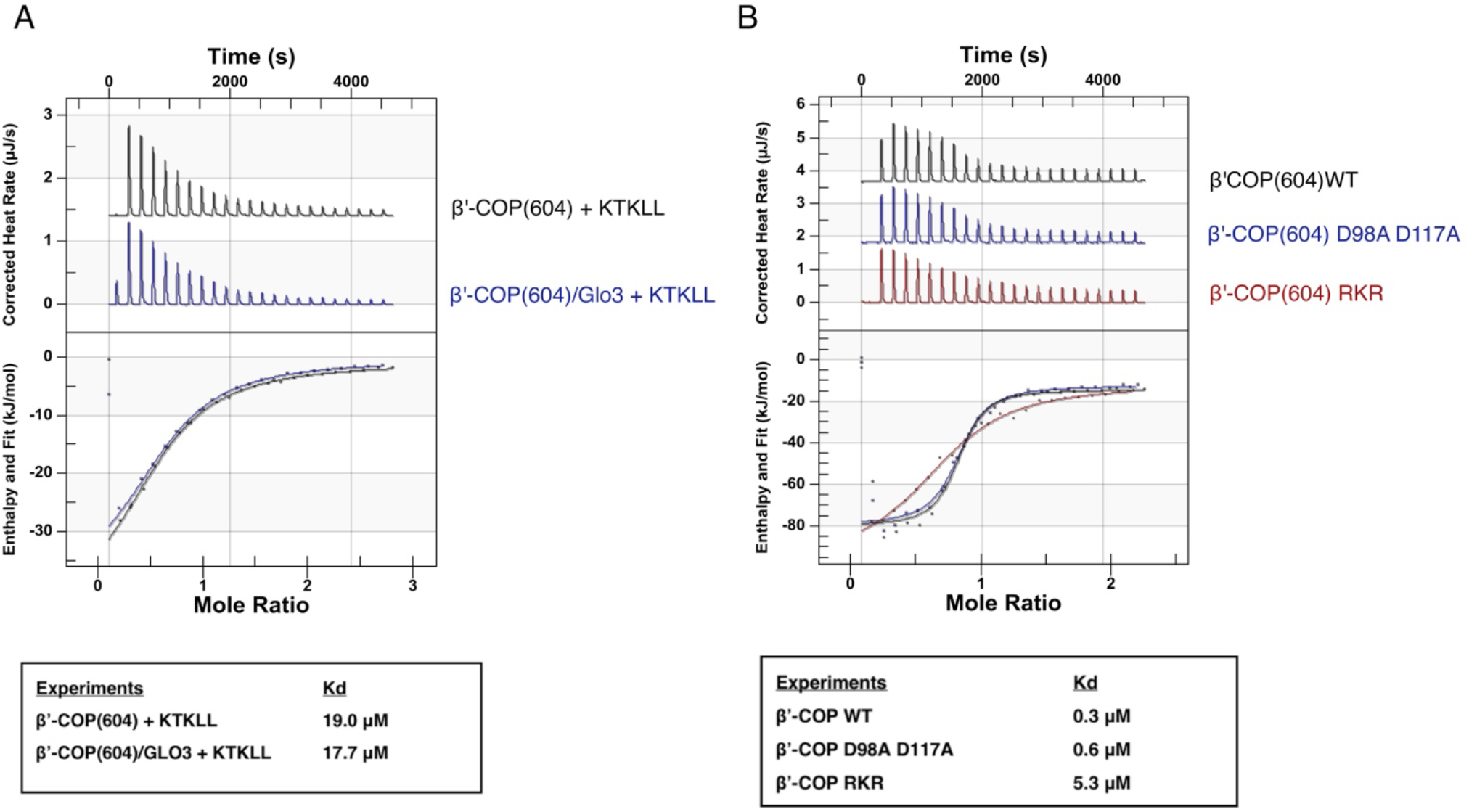
Glo3 BoCCS and dilysine cargo motifs bind β’-COP simultaneously *in vitro*. (A) Representative ITC experiments with untagged β’-COP 1-604; Glo3 residues 230-290; and dilysine motif (KTKLL) peptide. The presence of Glo3 does not alter the binding affinity of β’-COP to dilysine motifs *in vitro*. (B) ITC experiments with β’-COP dilysine binding mutants. The R13A/K15A/R59A (RKR mutant) disrupts binding to the dilysine motif carboxy-terminus, while D98A/D117A disrupts binding to lysine residues. The RKR mutant exhibits weaker binding to Glo3, while D98A/D117A binds Glo3 as well as wild-type protein. These data suggest Glo3 and dilysine motifs do not compete for binding β’-COP *in vitro*. However, disrupting the overall charge distribution on the N-terminal propeller in the RKR mutant suggests it may play some role in Glo3 binding.

**Figure S6.**
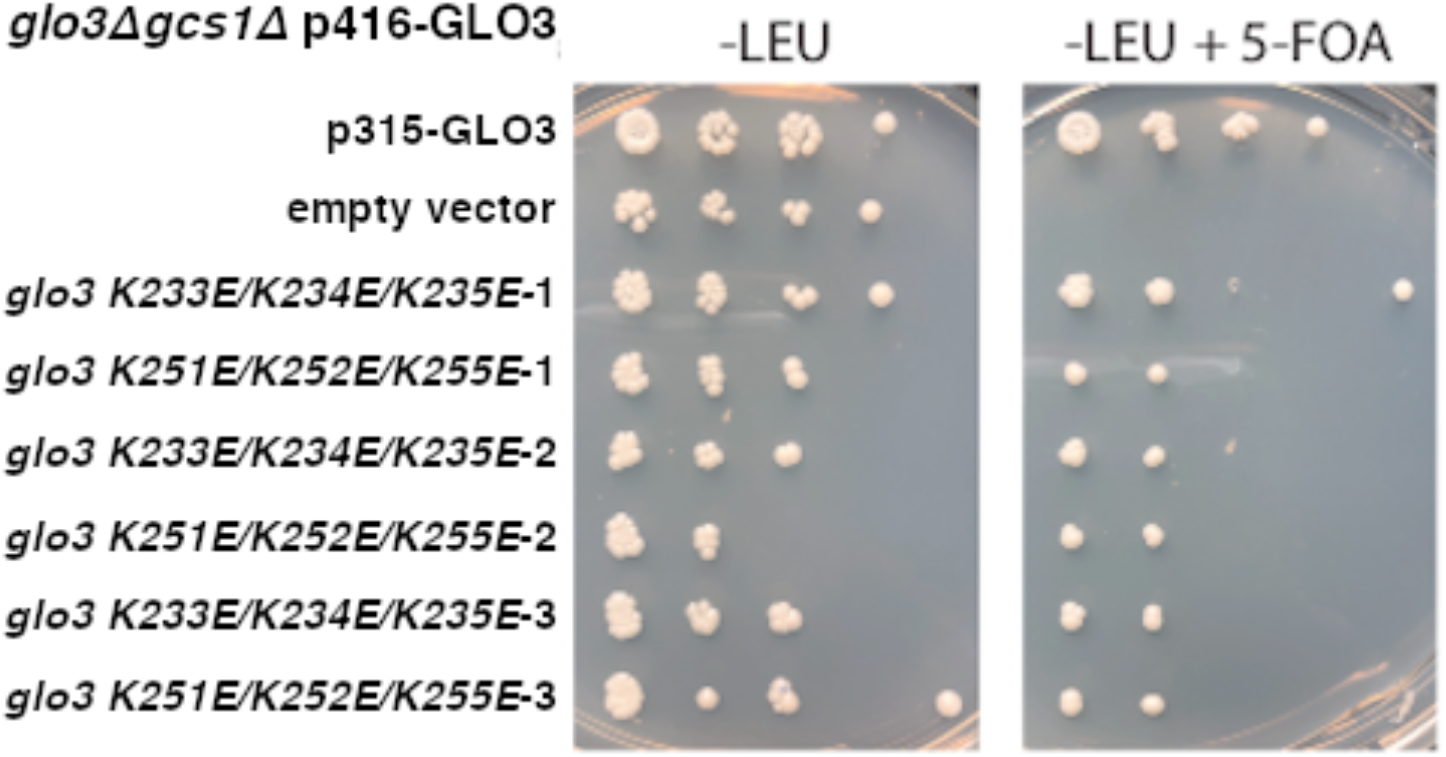
Yeast growth assays. Yeast growth assays in *glo3Δgcs1Δ* background strains with *GLO3* or *glo3* mutants (K233E/K234E/K235E or K251E/K252E/K255E). The *glo3Δ gcs1Δ* double mutant is inviable but can be sustained with a wild-type copy of *GLO3* on a *URA3* marked plasmid (pRS416). Both the wild type *GLO3* and the mutant forms were introduced into this background on a *LEU2* marked (pRS315) plasmid. On media that selects for both plasmids, all transformed strains grew like a wild-type strain because they contain the wild-type pRS416-*GLO3* plasmid. However, upon switching to 5-fluoro-orotic acid (5-FOA) media, which selects against the pRS416-*GLO3* plasmid, the *glo3Δ gcs1Δ* strain harboring the empty pRS415 plasmid failed to grow, as expected.

**Figure S7.**
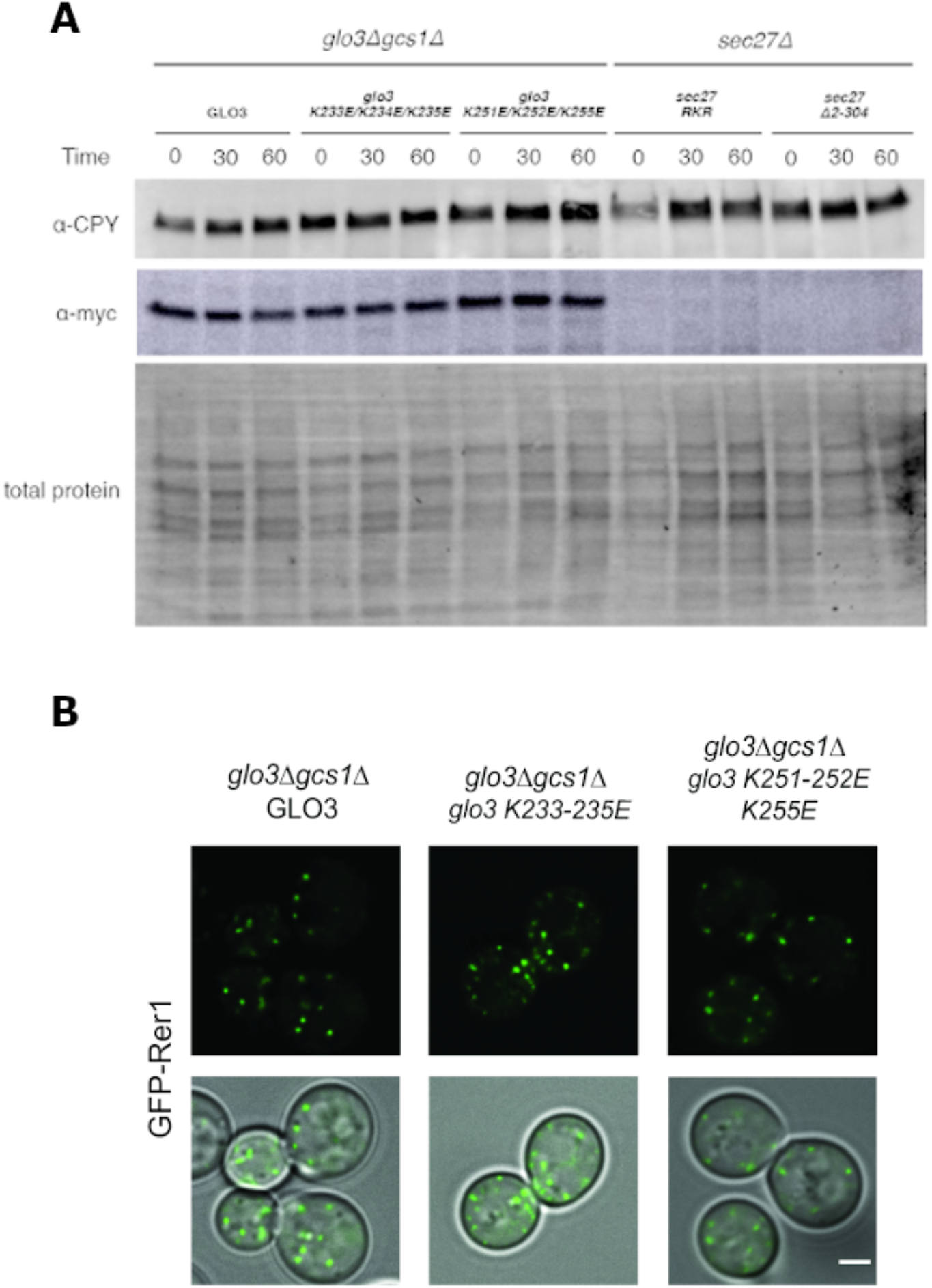
Golgi/ER COPI cargo trafficking data. (A) The Emp47-myc reporter construct (visualized using α−myc) remains stable over time in both wild-type *GLO3* and mutant *glo3* strains. In contrast, the reporter is sent to the vacuole and degraded when the dilysine binding site on β’-COP *(sec27)* is mutated (*sec27RKR;* R13A/K15A/R59A) or the first *sec27* propeller is deleted (*sec27Δ2-304*). (B) Representative fluorescence images of GFP-Rer1 in *glo3Δgcs1Δ* strains with *GLO3* or *glo3* mutants introduced on a plasmid. GFP-Rer1 trafficking does not appear to change when the β’-COP/Glo3 interaction is disrupted.

**Figure S8.**
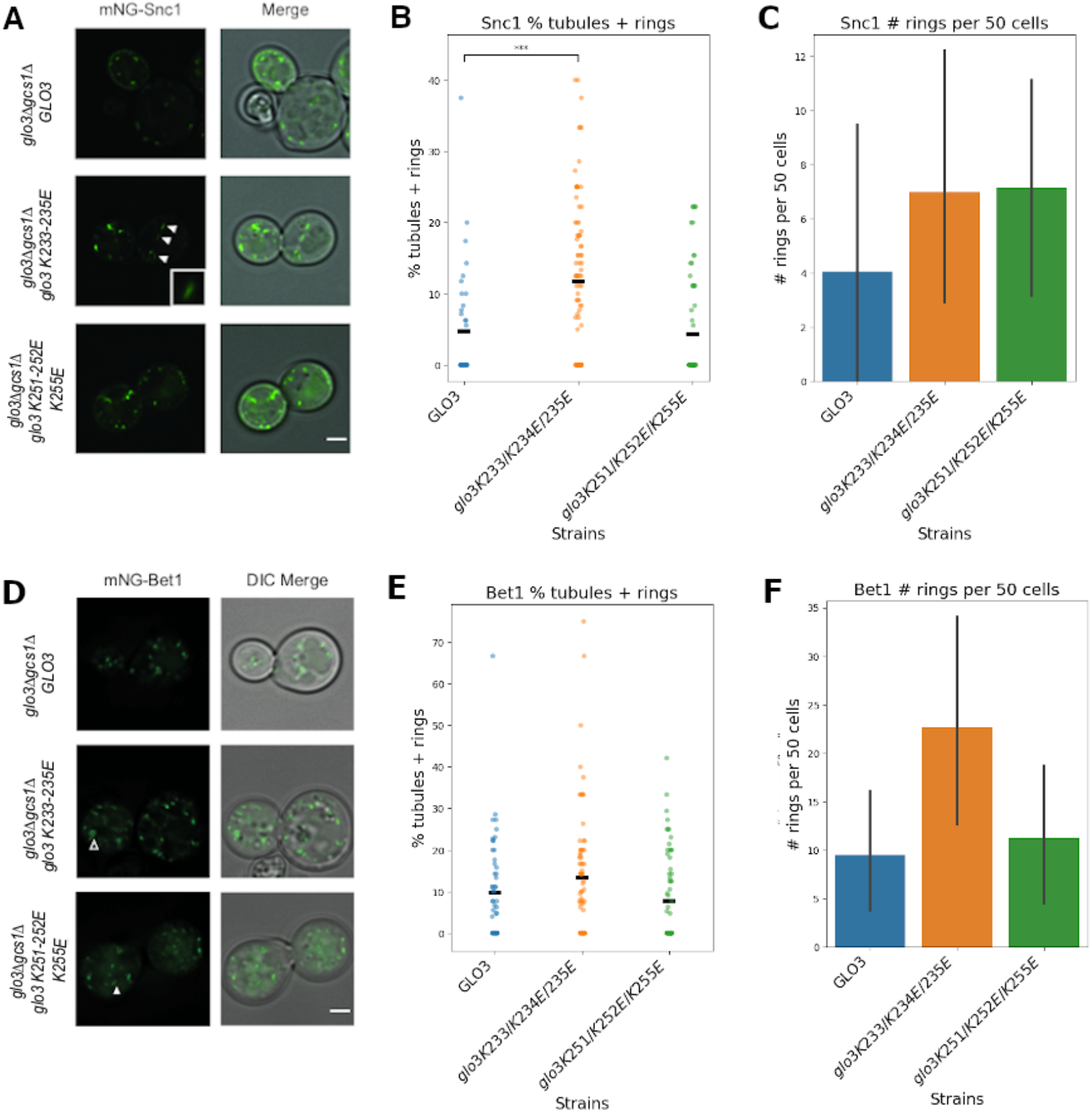
SNARE localization and Golgi morphology data. (A) Fluorescence imaging of mNG-Snc1 in *glo3Δ gcs1Δ* strains with *GLO3* or *glo3* mutants (K233E/K234E/K235E or K251E/K252E/K255E) on a plasmid. (B) Box plot showing percentage of abnormal structures (tubules and rings) with mean (black bar) observed in each strain. Mutating the first lysine cluster resulted in cells exhibiting a significant difference from wild-type. Significance was determined using a Mann-Whitney test comparing wild-type and mutant strains. (C) Histogram showing number of mNG-Snc1 ring structures with standard deviation (black lines) in each strain. (D) Fluorescence imaging of mNG-Bet1 in *glo3Δgcs1Δ* strains with *GLO3* or *glo3* mutants (K233E/K234E/K235E or K251E/K252E/K255E) on a plasmid. (E) Box plot showing percentage of abnormal structures (tubules and rings) with standard deviation observed in each strain. Although we sometimes observe abnormal structures, we do not find a significant difference, as determined by a Mann-Whitney test. (F) Histogram showing number of mNG-Bet1 ring structures with standard deviation (black lines) in each strain.

