## Supplementary material for "An interaction between β’-COP and the ArfGAP, Glo3, maintains post-Golgi cargo recycling": Table S1

| β'-COP | Glo3 | $K_D$ | n | Dilysine peptide | Trace shown in |
|---|---|---|---|---|---|
| 1-604 WT | 230-260 WT | 5.8 | 2.5 | -- | Figure 2 |
| 1-604 WT | 230-270 WT | 2.4 | 0.8 | -- | Figure 2 |
| 1-604 WT | 230-280 WT | 2.0 | 0.8 | -- | Figure 2 |
| 1-604 WT | 230-290 WT | 0.3 | 0.5 | -- | Figure 2, 3 |
| 1-604 WT | 230-300 WT | 0.8 | 0.8 | -- | Figure 2 |
| 1-304 WT | 230-300 WT | n.b. | n.b. | -- | Figure 2 |
| 300-604 WT | 230-300 WT | n.b. | n.b. | -- | Figure 2 |
| 1-604 WT | 208-383 WT | 0.8 | 0.8 | -- | Figure S3 |
| 1-604 WT | 220-300 WT | 0.8 | 0.7 | -- | Figure S3 |
| 1-604 WT | 230-300 WT | 0.8 | 0.8 | -- | Figure S3 |
| 1-604 WT | 230-290 K233E | 12 | 1.2 | -- | Figure 3 |
| 1-604 WT | 230-290 K233E/K234E/K235E | n.b. | n.b. | -- | Figure 3 |
| 1-604 WT | 230-290 251E/K252E/K255E | 12 | 0.9 | -- | Figure 3 |
| 1-604 D437A/D450A | 230-290 WT | 17.7 | 0.4 | -- | Figure 3 |
| 1-604 WT | 230-290 WT | 18.0 | 0.6 | KTKLL | Figure S5 |
| 1-604 R15A/K17A/R59A | 230-290 WT | 5.3 | 0.8 | -- | Figure S5 |
| 1-604 D98A/D117A | 230-290 WT | 0.6 | 0.8 | -- | Figure S5 |

**Table S1. ITC data summary.** This table summarizes representative ITC experiments, including protein constructs with residue numbers and relevant point mutations; calculated $K_D$ values; calculated stoichiometry (n) values; the addition of a dilysine peptide; and figure(s) showing relevant trace(s). "WT" denotes wild-type sequence.
