## Supplementary material for "An interaction between β’-COP and the ArfGAP, Glo3, maintains post-Golgi cargo recycling": Table S3

| β'-COP residues | Glo3 residues | Dilysine motif | Summary of results |
|---|---|---|---|
| 1-304 | None | KxKTN | Models do not converge or place dilysine motif in experimentally determined binding site |
| 1-304 | 230-290 | None | Models do not converge; Glo3 placed in multiple conformations around β'-COP periphery |
| 1-304 | 230-240 | None | Models do not converge; Glo3 K234 placed near β'-COP D98/D117 dilysine patch in some models |
| 1-304 | 250-260 | None | Models do not converge; Glo3 K252 or K255 placed near β'-COP D98/D117 dilysine patch in some models |
| 1-604 | None | KxKTN | Dilysine motif placed adjacent to C-terminal propeller instead of in experimentally determined binding site |
| 1-604 | 1-350 | None | Models do not converge; Glo3 K235 placed near β'-COP D523 in one model |
| 1-604 | 230-290 | None | Glo3 K286 residue placed near β'-COP D98/D117 in 3/5 models |
| 300-604 | 230-240 | None | Models do not converge |
| 300-604 | 250-260 | None | Models do not converge |

**Table S3. Summary of yeast β'-COP/Glo3 computational modeling experiments using AlphaFold2.**
