## Supplementary material for "An interaction between β’-COP and the ArfGAP, Glo3, maintains post-Golgi cargo recycling": Table S4

| **Yeast Strains** |
|---|
| *gcs1*Δ::kan *glo3*Δ::kan pRS315-GLO3 |
| *gcs1*Δ::kan *glo3*Δ::kan pRS315-*glo3 K233-235E* |
| *gcs1*Δ::kan *glo3*Δ::kan pRS315-*glo3 K251-252E, K255E* |
| *gcs1*Δ::kan *glo3*Δ::kan pRS315-GLO3 pRS416-mNG-Bet1 |
| *gcs1*Δ::kan *glo3*Δ::kan pRS315-*glo3 K233-235E* pRS416-mNG-Bet1 |
| *gcs1*Δ::kan *glo3*Δ::kan pRS315-*glo3 K251-252E, K255E* pRS426-mNG-Bet1 |
| *gcs1*Δ::kan *glo3*Δ::kan pRS315-GLO3 pRS416-mNG-Gos1 |
| *gcs1*Δ::kan *glo3*Δ::kan pRS315-*glo3 K233-235E* pRS416-mNG-Gos1 |
| *gcs1*Δ::kan *glo3*Δ::kan pRS315-*glo3 K251-252E, K255E* pRS416-mNG-Gos1 |
| *gcs1*Δ::kan *glo3*Δ::kan pRS315-GLO3 pRS416-GFP-Rer1 |
| *gcs1*Δ::kan *glo3*Δ::kan pRS315-*glo3 K233-235E* pRS416-GFP-Rer1 |
| *gcs1*Δ::kan *glo3*Δ::kan pRS315-*glo3 K251-252E, K255E* pRS416-GFP-Rer1 |
| *gcs1*Δ::kan *glo3*Δ::kan pRS315-GLO3 pRS416-mNG-Snc1 |
| *gcs1*Δ::kan *glo3*Δ::kan pRS315-*glo3 K233-235E* pRS416-mNG-Snc1 |
| *gcs1*Δ::kan *glo3*Δ::kan pRS315-*glo3 K251-252E, K255E* pRS416-mNG-Snc1 |
| *gcs1*Δ::kan *glo3*Δ::kan pRS315-GLO3 pRS426-GFP-Ste2 |
| *gcs1*Δ::kan *glo3*Δ::kan pRS315-*glo3 K233-235E* pRS426-GFP-Ste2 |
| *gcs1*Δ::kan *glo3*Δ::kan pRS315-*glo3 K251-252E, K255E* pRS426-GFP-Ste2 |

**Table S4. Yeast strains used in this study.**
